# Dynamics of macrophage polarization support *Salmonella* persistence in a whole living organism

**DOI:** 10.1101/2023.05.09.539693

**Authors:** Jade Leiba, Tamara Sipka, Christina Begon-Pescia, Matteo Bernardello, Sofiane Tairi, Lionello Bossi, Anne-Alicia Gonzalez, Xavier Mialhe, Emilio J. Gualda, Pablo Loza-Alvarez, Anne Blanc-Potard, Georges Lutfalla, Mai Nguyen-Chi

## Abstract

Numerous intracellular bacterial pathogens interfere with macrophage function, including macrophage polarization, to establish a niche and persist. However, the spatiotemporal dynamics of macrophage polarization during infection within host remain to be investigated. Here, we implement a model of persistent *Salmonella* Typhimurium infection in zebrafish, which allows visualization of polarized macrophages and bacteria in real time at high-resolution. While macrophages polarize toward M1-like phenotype to control early infection, during later stages, *Salmonella* persists inside non-inflammatory clustered macrophages. Transcriptomic profiling of macrophages confirmed a highly dynamic signature during infection characterized by a switch from pro-inflammatory to anti-inflammatory/pro-regenerative status and revealed a shift in adhesion program. In agreement with this specific adhesion signature, macrophage trajectory tracking identifies motionless macrophages as a permissive niche for persistent *Salmonella*. Our results demonstrate that zebrafish model provides a unique platform to explore, in a whole organism, the versatile nature of macrophage functional programs during bacterial acute and persistent infections.

## INTRODUCTION

The outcome of bacterial infections is the result of complex dynamic interactions between the pathogen and the host’s cellular and humoral actors of innate immunity. Deciphering this complexity is necessary to predict infection outcomes and guide therapeutic strategies. The development of tractable systems in which bacteria and cellular actors can be tracked at high spatio-temporal resolution in a whole living animal is essential to assess the dynamic of host-pathogen interactions.

*Salmonella enterica* is a gram-negative, facultative intracellular pathogen, inducing a variety of conditions ranging from benign gastroenteritis to severe systemic infection ^1^. Every year it infects 20 millions of people and cause more than 200,000 of deaths ^2^. In some cases, *Salmonella* establishes a chronic infection that results in asymptomatic carriers hosting the environmental reservoir for further infections ^3, 4^. In these asymptomatic carriers, *Salmonella* survives mainly inside macrophages where bacteria have to deal with this hostile environment ^5, 6^. Capitalizing on different virulence factors, *Salmonella* can replicate within macrophages inside modified vacuoles called phagosomes and escape the host’s defenses ^7^. In addition, as reported for several intracellular bacterial pathogens, *Salmonella* uses strategies to interfere with macrophage polarization ^8^.

Macrophages are among the most plastic immune cells that adapt their phenotype and function according to their microenvironment by a process called polarization. *In vivo*, they form a continuum of activation states whose two extremes are the pro-inflammatory M1 macrophages, that have a bactericidal activity, and the anti-inflammatory M2 macrophages that promote the resolution of inflammation and healing ^9^. During the first hours of infection, most bacteria, including *Salmonella enterica* serovar Typhimurium (*S.* Typhimurium), induce macrophage polarization toward the pro-inflammatory and microbicidal M1 phenotype ^10, 11^. By contrast, persisting bacteria, like *Mycobacterium tuberculosis*, *Brucella abortus* or *S.* Typhimurium, may preferentially reside inside healing/anti-inflammatory M2 macrophages ^5, 8, 12^. In a mouse model of *S.* Typhimurium long-term infection, bacteria were shown to persist mainly in M2 macrophages ^5, 13^. In contrast, another *in vivo* study showed that *S.* Typhimurium reside in iNOS-expressing macrophages that clustered within splenic granulomas 42 days post-infection in mice ^14^. Although iNOS is a known marker of M1 macrophages, the exact polarization status of these macrophages remained undetermined. In addition, intracellular replication of *S.* Typhimurium was shown to vary according to macrophage polarization, with a greater replication in M2 macrophages ^15^. To date, however, the dynamics of interactions between pathogenic bacteria and polarized macrophages between early invasion and late survival remain poorly understood at the organism level.

Perfectly suited to live observation of immune cells and pathogens *in vivo*, the transparent zebrafish embryo has emerged as a powerful vertebrate model to study host-pathogen interactions at the cellular and whole organism level ^16^. This model allows monitoring macrophage plasticity and reprogramming using dedicated fluorescent reporters to simultaneously tract macrophages and visualize their activation state. Using a non-infected wound model, we previously showed that macrophages first express pro-inflammatory cytokines (M1-like polarization) before switching to a new state expressing both M1 and M2 makers during the wound healing process ^17^. Yet, the dynamics of macrophage polarization in the context of *S.* Typhimurium infection has not been addressed in zebrafish.

Here we established the first larval zebrafish model of *S.* Typhimurium persistent infection. Using intravital imaging, we show that during early stages of infection, both neutrophils and macrophages are mobilized to the infection site to engulf bacteria and that macrophages respond by a strong M1-like activation. In later stages of infection, bacteria survive inside non-inflammatory macrophages which accumulate in large clusters and provide a niche for persistent bacteria inside the host. Finally, a comprehensive analysis of the transcriptional profiles of macrophages revealed a highly dynamic transcriptional signature of distinct macrophage subsets during early and late phases of infection, showing changes in inflammatory response and in expression of adhesion molecules upon persistent infection, which correlate with a decrease of macrophage motility. This new model provides a unique opportunity to explore the dynamics of interactions between persistent pathogenic bacteria and polarized macrophages in a 4-dimentional living system.

## RESULTS

### *Salmonella* HBV-infection in Zebrafish leads to different outcomes, from systemic to persistent infection

Previous studies on *S.* Typhimurium-infected zebrafish larva, based on intravenous or subcutaneous injection or immersion, have been linked to a rapid disease progression with acute symptoms and high production of pro-inflammatory cytokines, leading to larval mortality ^18, 19^. To develop a model of persistent bacterial infection using *S.* Typhimurium (hereafter named *Salmonella)*, we chose to inject in a closed compartment, the HindBrain Ventricle (HBV) (**Fig. 1A**), which allows direct visualization of immune cell recruitment in a confined space. To allow direct observation of bacterial burden, a GFP-expressing *Salmonella* ATCC14028s strain was constructed (Sal-GFP). We injected 2 days post-fertilization (dpf) zebrafish embryos with different doses of Sal-GFP in the HBV, or with PBS as control (**Fig. 1B**) and monitored larval survival from 0 to 4 days post-infection (dpi). Increasing injection doses of *Salmonella* resulted in increased larval mortality. When less than 500 Colony Forming Units (CFU) of Sal-GFP were injected, all zebrafish larvae survived the infection (**Fig. 1C**). On the other hand, HBV injection of 1000 to 2000 CFU of Sal-GFP led to 50% of larvae survival at 4 dpi (**Fig. 1D**).

**Fig. 1.**
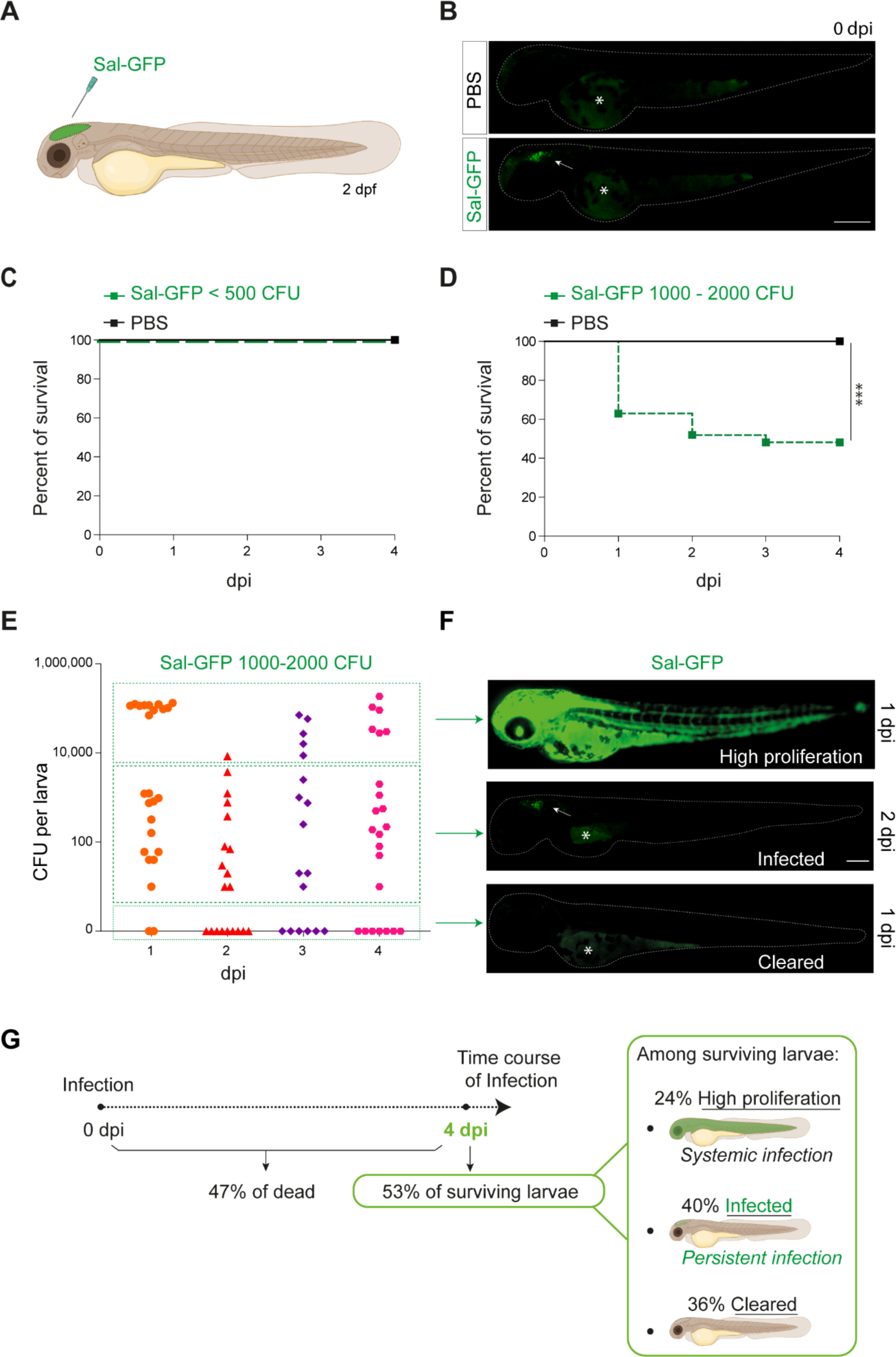
Zebrafish is a pertinent model for persistent *Salmonella* infection. **(A)** Schematic illustration of 2 dpf zebrafish embryo infected in the Hind-brain Ventricle (HBV) with Sal-GFP, a GFP-expressing strain. **(B)** Representative fluorescent images of HBV-injected larvae with either PBS or 1000 CFU of Sal-GFP shortly after micro-injection. White arrow: bacteria in the HBV. Dots outline the larva. Asterisk: auto-fluorescence of the yolk. Scale bar: 200 μm. **(C-D)** Survival curves of injected embryos with either PBS or different doses of Sal-GFP, i.e. **(C)** < 500 CFU or **(D)** 1000 – 2000 CFU. One representative of three replicates (n=24 larvae per condition). Log rank test, *** *P* < 0.001. **(E)** CFU counts per embryos infected with a range of 1000 to 2000 CFU of Sal-GFP at 1, 2, 3 and 4 dpi. Pool of four independent experiments (n_1dpi_=25, n_2dpi_=20, n_3dpi_=20, n_4dpi_=25 larvae). **(F)** Representative fluorescent images of Sal-GFP infected larvae. Bacteria are in green. Dots outline the larva. Asterisk: auto-fluorescence of the yolk. Scale bar: 200 μm. **(G)** Schematic representation of the different infection outcomes, **High Proliferation**, **Infected** and **Cleared**, induced by injection of 1000 – 2000 CFU of Sal-GFP. From 0 to 4 dpi, 47% of the infected larvae developed a systemic infection where the bacteria displayed highly-proliferation leading to larval death (**High Proliferation)**. At 4 dpi, among the surviving larvae, 24% still exhibited a systemic infection, while 36% recovered from the infection with no detectable CFU (**Cleared)** and 40% contained persistent bacteria **(Infected)**.

To evaluate the bacterial burden in infected hosts, the number of CFU were counted every day from 0 to 4 dpi after HBV-injection of Sal-GFP (**Fig. 1E**). Before processing for CFU counting, infected larvae and their controls were imaged individually by fluorescence microscopy. No CFU were detected in the PBS-injected larvae (**Fig. S1A**). After injecting 1000-2000 CFU, bacteria efficiently proliferated in 47% of infected larvae, from 1 to 4 dpi, (50 000-200 000 CFU, **Fig. 1E** and **1G**). Those larvae developed a systemic infection with *Salmonella* proliferation in the HBV and other tissues, including notochord, heart and circulating blood (**Fig. 1F**) and usually died between 2 and 4 dpi. Among the 53% of infected larvae that survived the infection at 4 dpi, 40% harbored live bacteria (10 to 2000 CFU), while 36% were bacteria-free (**Fig. 1E and 1G**). When injecting less than 500 CFU, no larva exhibited bacterial hyper-proliferation (**Fig. S1B**) and 75% showed evidence of *Salmonella* persistence at 4 dpi (10 to 1000 CFU, **Fig. S1B**). Importantly, alive infected larvae still harbored persistent bacteria at 14 dpi (**Fig. S1C**).

Altogether, these results show that *Salmonella* infection in HBV leads to different outcomes, that we classified as: i, **High Proliferation**, indicating a high bacterial burden with systemic infection leading to death; ii, **Infected**, indicating surviving larvae with persisting bacteria; and iii, **Cleared**, indicating that larvae completely recovered with no more detectable bacteria (**Fig. 1G**).

Throughout the remainder of this study, an optimized infection dose of 1000 to 1500 CFU was injected into the HBV of 2 dpf embryos and we further focused (unless otherwise stated) on the so called “infected” cohort with established persistent infection.

### The global host inflammatory response to *Salmonella* HBV-infection

To investigate the global host immune response to *Salmonella* infection in the zebrafish model, the relative expression of several immune-related genes was examined by qRT-PCR within whole larvae from 3 hpi to 4 dpi after *Salmonella* challenge (**Fig. 2** and **Fig. S1D**). At 3 hpi, infected larvae showed elevated levels of expression of pro-inflammatory cytokines: *interleukin-1 beta* (*il-1b)*, *tumor necrosis factor a* and *b* (*tnfa* and *tnfb*, two orthologs of mammalian *TNF*), *interleukin-8* (*il-8*) and of the inflammation marker *matrix metalloproteinase 9* (*mmp9*) (**Fig. 2A**), consistent with previous findings in zebrafish models of systemic *Salmonella* infection^18^. Subsequently, pro-inflammatory gene expression (*il-1b*, *tnfb*, *il-8* and *mmp9*) significantly decreased at 1 dpi and raised from 2 hpi to 4 dpi to reach similar levels to those detected at 3 hpi. At 4 dpi, *tnfa* expression was still up-regulated in infected larvae compared to PBS controls (**Fig. S1D**). *Salmonella* infection also induced *ccl38a.4* gene, encoding CCL2 a key chemokine of macrophage migration that binds the CCR2 receptor ^20^, suggesting a role of *ccl38a.4* in macrophage mobilization to the infection site. The up-regulation of the macrophage specific marker *mfap4* during the time course of infection highlighted an overall macrophage response (**Fig. 2A**). In contrast, we found that the other frequently used macrophage specific marker *mpeg1* was down-regulated early after *Salmonella* infection (**Fig. 2A**), as previously described during *Salmonella* and *Mycobacterium marinum* infections in zebrafish^21^. Then, the kinetics of expression of anti-inflammatory genes revealed a down-regulation of *mannose receptor, C type 1b* (*mrc1b*) early after *Salmonella* infection. Intriguingly, relative levels of *mrc1b* expression increased from 3 hpi to 3 dpi. Furthermore, *Negative Regulator of Reactive Oxygen Species* (referred to as *nrros*), regulator of reactive oxygen species and of TGF-b in mammals ^22–25^, showed kinetic of expression similar to *mrc1b* during *Salmonella* infection.

**Fig. 2.**
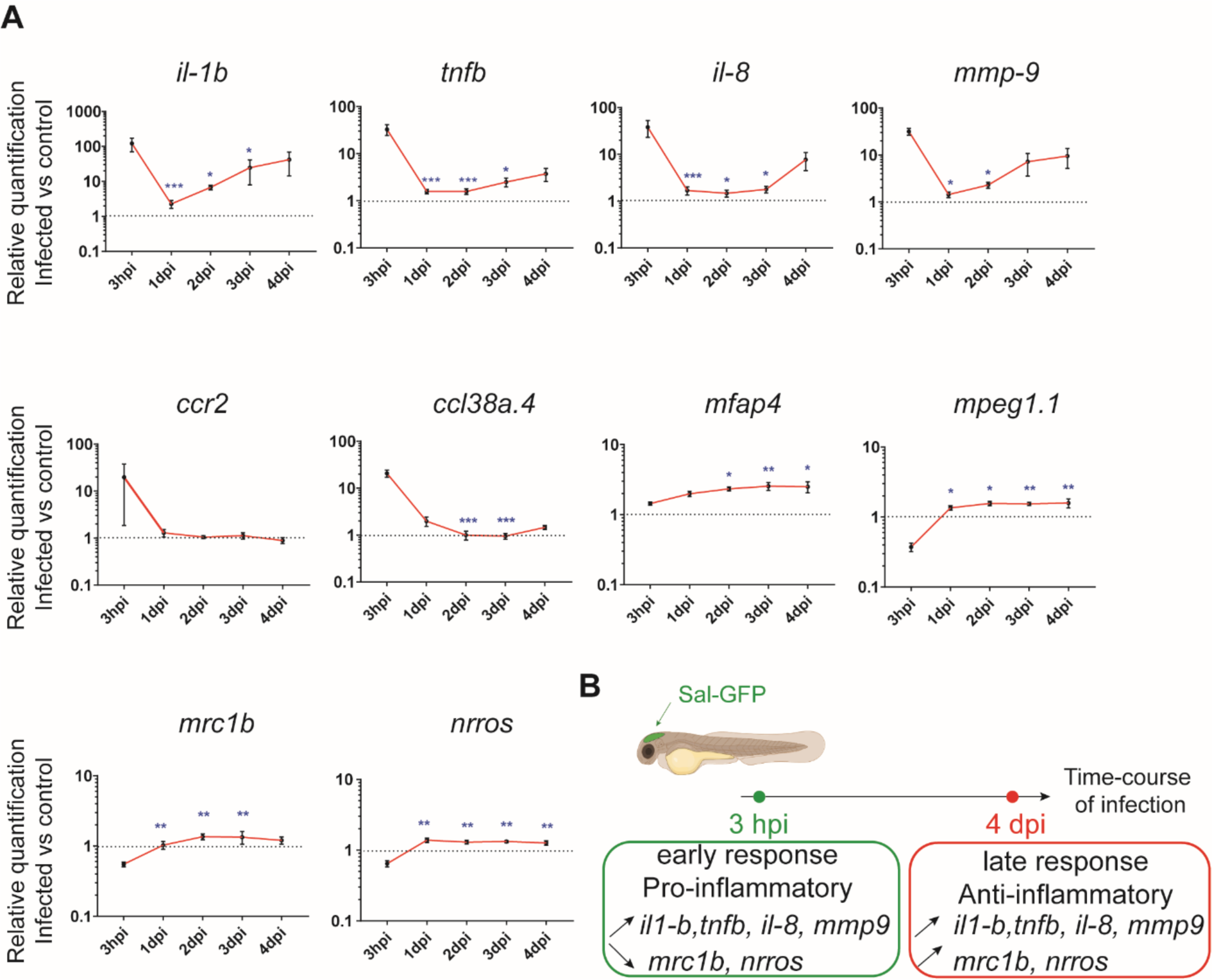
The global host inflammatory response to *Salmonella* infection. **(A)** qPCR analysis of *il1b*, *tnfb*, *il8*, *mmp9*, *ccr2*, *ccl38a.4*, *mpeg1*, *mfap4, mrc1b* and *nrros*, mRNA expression infected *vs* non-infected, normalized with *ef1a.* Larvae were either PBS- or Sal-GFP injected and RNA samples were extracted from whole larvae at 3 hpi, 1, 2, 3and 4 dpi. Data are presented as relative expression in the infected larvae compared with the relevant PBS-injected controls (2^-1¢1¢Cp^). Values are the means ± SEM of eight replicates (n= 8 larvae per time point). Kruskal-Wallis test (unpaired, non-parametric) * *P* < 0.05; ** *P* < 0.01; *** *P* < 0.001 show significant differences compared to 3 hpi. **(B)** Diagram of global host inflammatory response to *Salmonella* infection.

Taken together these results highlight a specific immune response triggered by *Salmonella*, and define two main phases: an early pro-inflammatory phase characterized by the induction of pro-inflammatory markers and down-regulation of anti-inflammatory markers, and a late phase (between 3 and 4 dpi) characterized by *de novo* up-regulation of pro-inflammatory gene expression concomitant with the increased expression of anti-inflammatory genes (**Fig. 2B**). Based on these results, we decided to further focus our study on deciphering the immune response to *Salmonella* infection both during the early phase, few hours post-infection, and during the late phase at 4 dpi.

### Early phase of *Salmonella* HBV-infection induces strong macrophage and neutrophil responses

Among immune cells, macrophages are a niche for *Salmonella* infection in mammals ^1, 6^ and in zebrafish ^18, 19^. To investigate the role of macrophages in the control of *Salmonella* infection, macrophage reporter embryos, Tg(*mfap4:mCherry-F*) were injected in the HBV with Sal-GFP or PBS and the global macrophage population was imaged by fluorescence microscopy from 3 hpi to 4 dpi (**Fig. 3A**). We noticed that *Salmonella* infection led to an increase in the global macrophage population that persisted over days compared to PBS-injected larvae, suggesting that *Salmonella* infection triggers myelopoiesis (**Fig. 3A-B**). At 3 hpi macrophages were specifically recruited to the HBV of infected embryos. In addition, at 4 dpi, an intense macrophage accumulation persisted in the brain of infected embryos, in line with the development of a *Salmonella* persistent localized infection. To test whether *Salmonella* persistence depends on the infection site, we injected either Sal-GFP or PBS in the otic vesicle of 2 dpf macrophage reporter embryos, Tg(*mfap4:mCherry-F*) (**Fig. S1E**). At 4 dpi, intravital confocal imaging revealed macrophage recruitment to the infected otic vesicle and the presence of bacteria, confirming the intrinsic capacity of *Salmonella* to persist within the host for a long time (**Fig. S1F**).

**Fig. 3.**
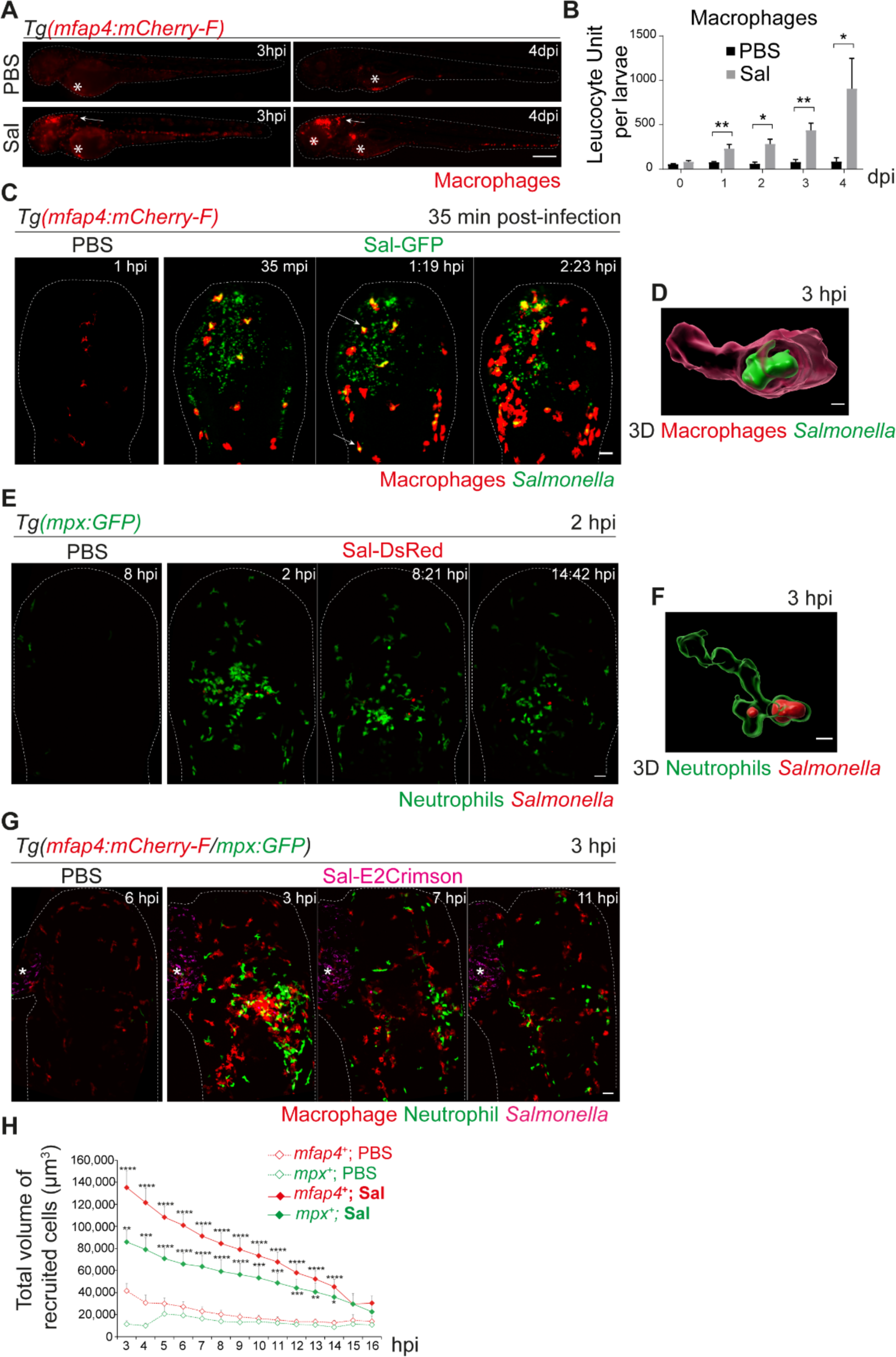
Early phase of *Salmonella* HBV-infection induces strong macrophage and neutrophil responses. **(A-D)** Tg*(mfap4:mCherry-F)* larvae were injected with either PBS or Sal-GFP in HBV. **(A)** Representative fluorescent images of larvae showing macrophage recruitment at the site of injection at 3 hpi and at 4 dpi. Asterisk: auto-fluorescence. Scale bar: 200 μm. **(B)** Quantification of total macrophages at 0, 1, 2, 3 and 4 dpi. One representative of three replicates. (Mean number of Leukocyte Unit/larva ± SEM, n_0dpi_= 24, n_1dpi_= 11, n=_2,3,4dpi_= 5 per condition, Mann-Whitney test, two-tailed, * *P* <0.05, ** *P* <0.01). **(C)** Representative maximum projections of fluorescent images extracted from 4D sequences using light sheet fluorescence microscopy starting 35 min post-infection during 2 hours, showing recruitment of macrophages (red) to the infection site (*Salmonella*, green). Scale bar: 30 μm. **(D)** 3D reconstruction of a macrophage phagocytosing *Salmonella* at 3 hpi. Scale bar: 5 μm. **(E-F)** Tg*(mpx:GFP)* larvae were injected with PBS or Sal-DsRed in HBV. **(E)** Representative maximum projections of fluorescent images extracted from 4D sequences using confocal microscopy at 2 hpi during 13 hours, showing recruitment of neutrophils (green) to the infection site (*Salmonella*, red). Scale bar: 35 μm. **(F)** 3D reconstruction of a neutrophil phagocytosing *Salmonella* at 2 hpi. Scale bar: 5 μm. **(G-H)** Tg*(mfap4:mCherry-F/mpx:GFP)* larvae were injected with either PBS or Sal-E2Crimson in HBV. **(G)** Representative maximum projections extracted from 4D sequences using confocal microscopy from 3 to 14 hpi showing recruitment of both neutrophils (green) and macrophages (red) to the infection sites. Asterisk: auto-fluorescence. Scale bar: 50 μm. **(H)** Quantification of the total volume of recruited cells (*mfap4*^+^ or *mpx*^+^ cells) from 3 to 16 hpi. Data of three replicates pooled. (Mean volume/larva ± SEM, n= 15 per condition, Mann-Whitney test, two-tailed, significance of Sal versus PBS conditions * *P* <0.05, ** *P* <0.01, *** *P* <0.001, **** *P* <0.0001).

To obtain a 4D (space + time) description of macrophage interactions with *Salmonella*, we used a Light Sheet Fluorescence Microscopy (LSFM) system, convenient to study the 3D architecture of cells over a large field of view and long periods of time with minimal side effects. Tg(*mfap4:mCherry-F*) larvae were infected with Sal-GFP and imaged. The resulting video sequences revealed macrophage recruitment within 2 hpi (**Fig. 3C** and **Movie S1**) and immediate *Salmonella* internalization (**Fig. 3C**, arrowheads, **Fig. 3D** and **Movie S2**).

To investigate the role of macrophages in the control of *Salmonella* infection, we ablated macrophages using Tg(*mpeg1:Gal4/UAS:nfsB-mCherry*) embryos and metronidazole (MTZ) treatment ^26^ (**Fig. S2A-B**). MTZ (added at 48 hours post-fertilization (hpf)) specifically ablated 71% of macrophages in transgenic larvae after 24 h of treatment (**Fig. S2C-D**). DMSO treatment on transgenic larvae (NTR+MTZ-) and metronidazole treatment on WT embryos (NTR-MTZ+) were used as controls (**Fig. S2C-H** and not shown). PBS injection had no effect on the mortality of embryos from the different groups, confirming the absence of toxicity of MTZ treatment (**Fig. S2E**). After infection with a sublethal dose of 500 CFU of Sal-GFP, MTZ-mediated macrophage depletion increased larval mortality up to 54% at 4 dpi while control group showed 4% mortality (**Fig. S2F**). MTZ-mediated macrophage depletion also strongly impacted bacterial clearance with development of a systemic infection, as shown by fluorescence microscopy and quantification of bacterial burden (**Fig. S2G-H**). Altogether, these data show that macrophages are instrumental to control *Salmonella* infection in HBV.

Neutrophils are also innate immune players in the defense against *Salmonella* in various models ^27, 28^. To investigate their role in our model, we infected Tg(*mpx:GFP*) neutrophil reporter embryos with DsRed-fluorescent *Salmonella* (Sal-DsRed). The fluorescence analysis of global neutrophil populations showed an increase within 2 dpi, in line with the reported *Salmonella*-induced granulopoiesis ^28^ (data not shown). Time-lapse imaging was used to visualize early neutrophil-bacteria interactions (**Fig. 3E** and **Movie. S3**): while neutrophils were absent from PBS-injected HBV, they were immediately recruited in the infected HBV and participated to bacterial clearance through phagocytosis. Subsequently, after 3 hpi, the number of neutrophils decreased at the infection site (**Fig. 3E-F**).

To assess the relative contribution of both macrophage and neutrophil to control *Salmonella* infection, Tg(*mfap4:mCherry-F*/*mpx:GFP*) embryos were infected in the HBV with E2Crimson fluorescent *Salmonella* (Sal-E2Crimson) and imaged from 3 to 14 hpi (**Fig. 3G** and **Movie. S4**). Because counting individual leukocytes that stack together at the infection site is not reliable, we measured the total of volume of both macrophages (*mfap4*^+^ cells) and neutrophils (*mpx*^+^ cells) using 3D reconstructions. Total volume analysis showed that macrophages and neutrophils were both strongly recruited to the infection site in the first hours of infection compared to controls and that their mobilization slightly diminished over time post-infection with similar kinetics (**Fig. 3G-H**).

Altogether, we concluded that both macrophages and neutrophils actively participate to the immediate innate immune response to *Salmonella* infection in the HBV, characterized by myelopoiesis, granulopoiesis, leukocyte recruitment to the infection site and phagocytosis of bacteria.

### *Salmonella* HBV-infection induces hyper accumulation of macrophages harboring persistent bacteria at late stage of infection

To image long-term *Salmonella* infection, we used a *Salmonella* strain harboring a stable chromosomal version of the *GFP* gene (Sal-GFP-int). Macrophage reporter embryos Tg(*mfap4:mCherry-F*) were infected with Sal-GFP-int and imaged at 4 dpi using intravital confocal microscopy. Infected larvae displayed a massive accumulation of macrophages in the HBV, with clusters associated with persisting fluorescent *Salmonella* (**Fig. 4A**, shown with arrows). 3D reconstruction of *mfap4*-positive cell volumes from confocal images revealed that *Salmonella* resided mainly inside some macrophages and only few *Salmonella* outside macrophages were observed (**Fig. 4B**). Similar experiments with neutrophil reporter embryos Tg(*mpx:GFP*) showed that although persisting bacteria could be observed, few neutrophils were present in the HBV at 4 dpi, but they were not clustered and did not contain persistent *Salmonella* (**Fig. 4C-D**).

**Fig. 4.**
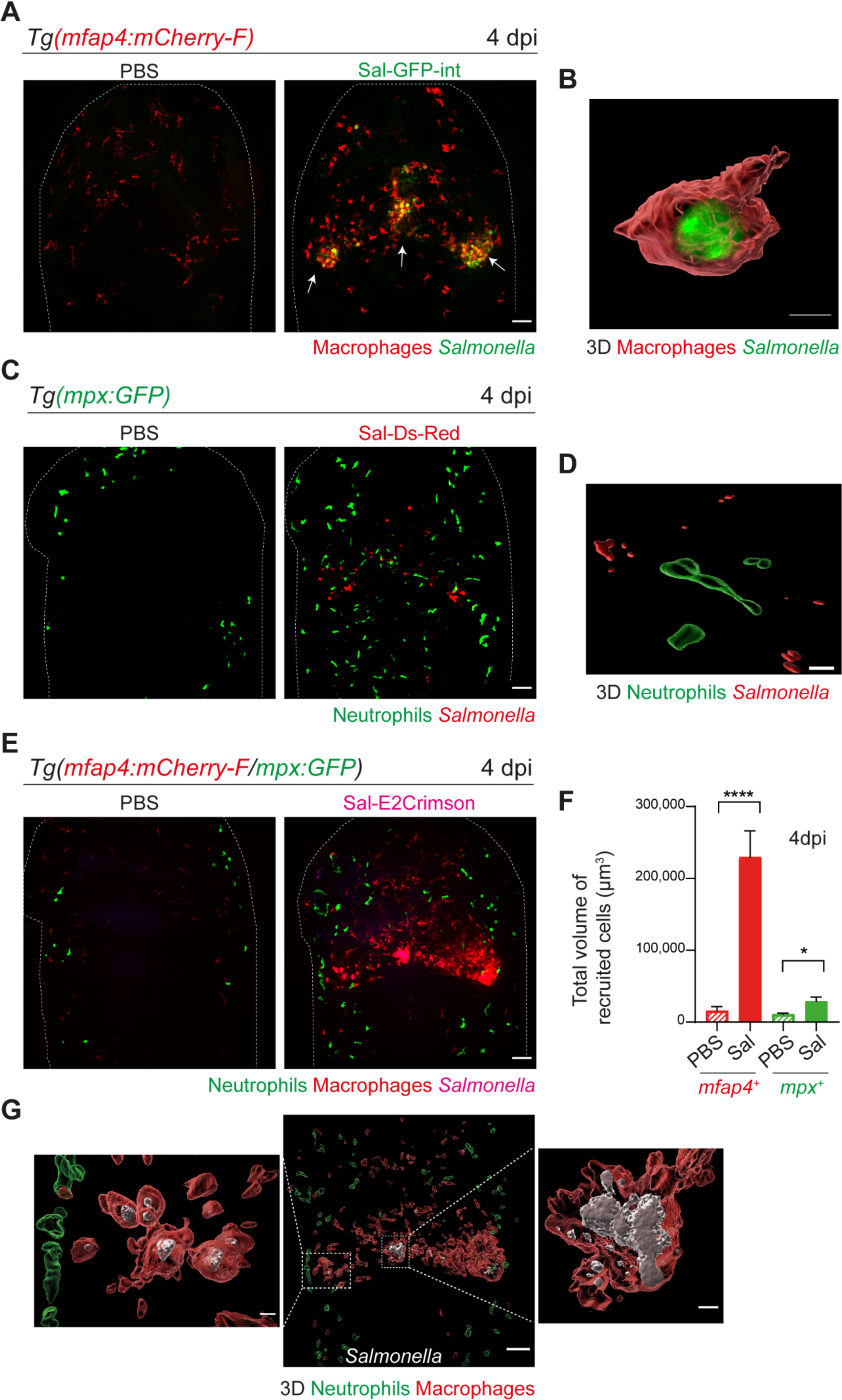
*Salmonella* HBV-infection induces hyper-accumulation of macrophages harboring persistent bacteria at late time point of infection. **(A)** Tg*(mfap4:mCherry-F)* larvae were injected with either PBS or Sal-GFP-int in HBV. Representative maximum projections of fluorescent confocal images, showing accumulation of macrophages (red) that co-localize with persistent *Salmonella* (green) at 4 dpi. Scale bar: 50 μm. **(B)** 3D reconstruction of persistent *Salmonella* residing inside a macrophage at 4 dpi. Scale bar: 5 μm. **(C)** Tg*(mpx:GFP)* larvae injected with either PBS or Sal-DsRed in HBV. Representative maximum projections of fluorescent confocal images showing that *Salmonella* (red) do not co-localize with neutrophils (green) at 4 dpi. Scale bar: 50 μm. **(D)** 3D reconstruction of neutrophils and persistent *Salmonella* at 4 dpi. Scale bar: 30 μm. **(E-G)** Tg*(mfap4:mCherry-F/mpx:GFP)* larvae were injected with either PBS or Sal-E2Crimson in HBV. **(E)** Representative maximum projections of fluorescent confocal images showing macrophage clusters (red), persistent *Salmonella* (magenta) and neutrophils (green) at 4 dpi. Scale bar: 50 μm. **(F)** Quantification of the total volume of recruited cells (*mfap4*^+^ or *mpx*^+^ cells) at 4 dpi. Data of three replicates pooled. (Mean volume/larva ± SEM, n_Sal_= 20, n_PBS_= 8, Mann-Whitney test, two-tailed, * *P* < 0.05, **** *P* < 0.0001). **(G)** 3D reconstruction of the HBV (middle panel) showing macrophage clusters (red) in which *Salmonella* (grey) persist, surrounded by neutrophils (green) at 4 dpi. Right and left panels are zooms of regions boxed by dotted lines. Scale bar: 30 μm. Scale bar zooms: 10 μm.

To simultaneously visualize the relative position of both macrophages and neutrophils at the infection site with persistent *Salmonella*, Tg(*mfap4:mCherry-F*/*mpx:GFP*) embryos were infected with Sal-E2Crimson and imaged at 4 dpi (**Fig. 4E** and **Movie. S5**). Quantification of the total volume of recruited macrophages (*mfap4*^+^ cells) and neutrophils (*mpx*^+^ cells) in HBV confirmed an important macrophage recruitment, which was five time stronger at 4 dpi than at 16 hpi (**Fig. 3H** and **Fig. 4F**). By contrast, neutrophils were similarly recruited at 4 dpi compared to 16 hpi. Importantly, this experiment confirmed that neutrophils do not co-localize with persistent *Salmonella* at late stages of infection (**Fig. 4G**).

These findings show that at later stages of infection, *Salmonella* infection triggers a differential macrophage and neutrophil response characterized by a massive macrophage recruitment in the HBV with formation of large macrophage clusters containing persistent bacteria.

### During *Salmonella* infection, macrophages first acquire a M1-like phenotype and polarize toward non-inflammatory phenotype at later stages

The initial macrophage response to *Salmonella* infection was previously shown to be pro-inflammatory in mouse and zebrafish models ^5, 13, 29–31^. We thus hypothesized that zebrafish macrophages polarize toward M1-like phenotype upon *Salmonella* infection. The pro-inflammatory cytokine TNFa is a known marker for M1-like macrophage in various species including zebrafish ^17^. First, the transgenic line Tg(*tnfa:GFP-F*) was used to track Tnfa producing cells by expression of a farnesylated GFP (GFP-F) during acute infection. Embryos were infected with Sal-DsRed and imaged every 5 min from 1 to 10 hpi. Time-lapse videos showed that GFP-F expression was induced at the infection site and labelled cells harbored a myeloid morphology, suggesting that they are macrophages (**Fig. S3A** and **Movie. S6**). To investigate the dynamic of macrophage polarization during *Salmonella* infection *in vivo*, we used the double transgenic line Tg(*mfap4:mCherry-F*/*tnfa:GFP-F*) in which all macrophages express a farnesylated mCherry and cells producing Tnfa, express GFP-F ^17^. Tg(*mfap4:mCherry-F*/*tnfa:GFP-F*) larvae were infected with Sal-E2Crimson and imaged at early time and at 4 dpi (**Fig. 5** and **Movie. S7**). Quantification of volumes of *mfap4*- and *tnfa*-positive cells, every hour from 4 to 15 hpi, showed that *Salmonella* infection induced a strong *tnfa* response increasing over time, unlike PBS-injected controls in which only few *tnfa*-positive cells were observed (**Fig. 5A-C** and **Fig. S3A**). During the first hours of infection, a maximum of 80% of *mfap4*-positive cells became also *tnfa*-positive (*mfap4*^+^*tnfa*^+^ cells) corresponding to M1-like activated macrophages (**Fig. 5B**). Interestingly both macrophages with and without bacteria expressed *tnfa* (**Fig. 5C**). These results show that in the early phase of *Salmonella* infection, most of the macrophages polarize towards M1-like, including both infected and bystander macrophages.

**Fig. 5.**
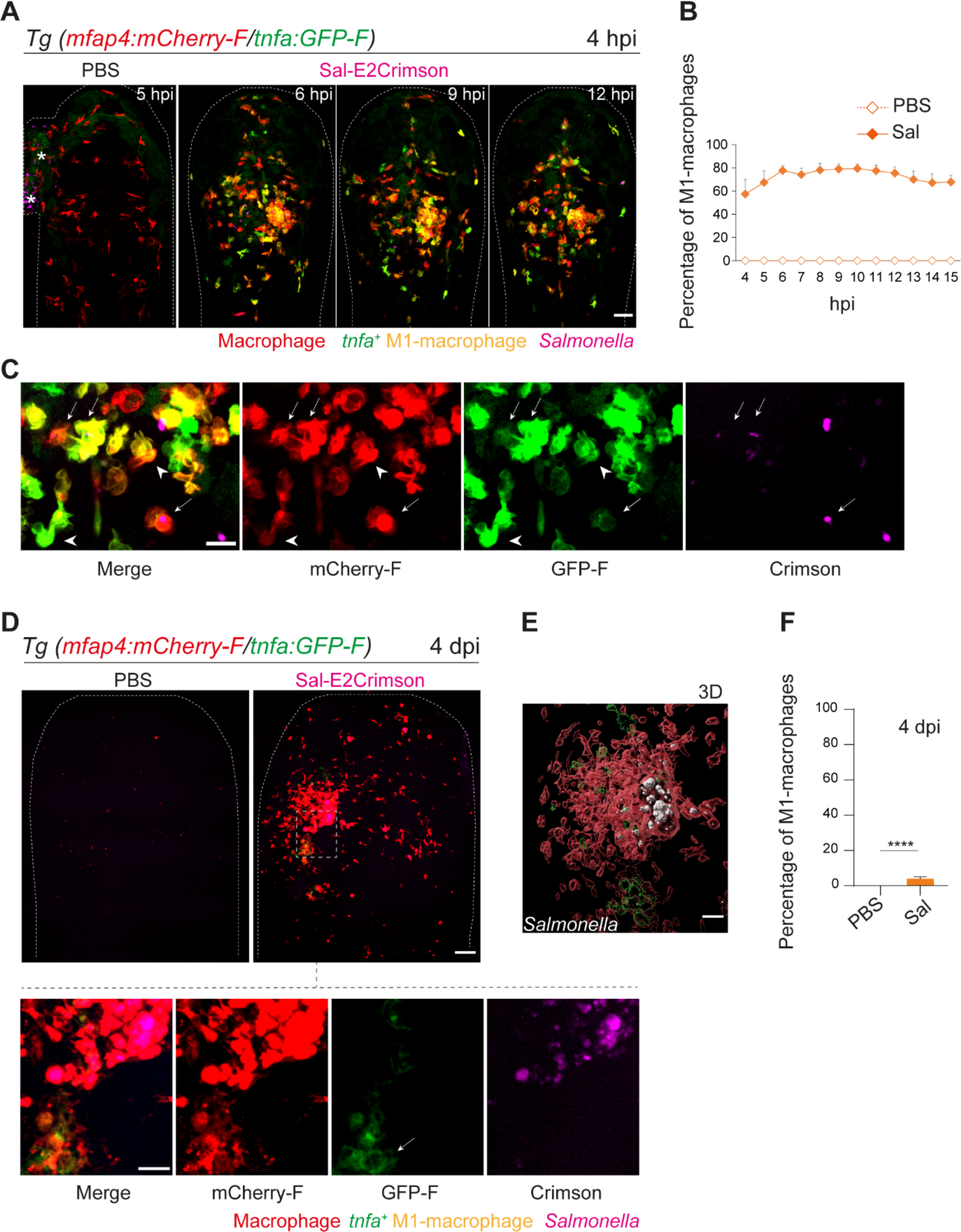
Macrophages polarize toward a pro-inflammatory M1-like phenotype upon *Salmonella* infection at early stage but not at late stage. **(A-F)** Tg*(mfap4:mCherry-F/tnfa:GFP-F)* larvae were injected with PBS or Sal-E2Crimson in HBV. **(A)** Representative maximum projections of fluorescent confocal images extracted from a 4D sequence, showing recruitment of macrophages (*mfap4*^+^ cells, red) and M1-like activation (*mfap4*^+^-*tnfa*^+^ cells, yellow) to *Salmonella* (magenta) from 4 to 15 hpi. Asterisk: auto-fluorescence. Scale bar: 50 μm. **(B)** Quantification of the percentage of M1-macrophages at indicated time post-infection. Data of two replicates pooled. (Mean percentage/larva ± SEM, n= 12 per condition). **(C)** Zoom of fluorescent confocal images in A. Scale bar: 20 μm, arrow: infected *tnfa*^+^ macrophage and arrowhead *tnfa*^+^ bystander macrophage. **(D)** Representative maximum projections of fluorescent confocal images of PBS-injected and Sal-E2Crimson infected larvae at 4 dpi (upper panels). Scale bar: 50 μm. Zooms of regions boxed by dotted lines (bottom panels). Scale bar zoom: 10 μm. **(E)** 3D reconstruction of macrophage clusters (red) containing persistent *Salmonella* (grey), surrounded by few *tnfa*^+^ macrophages (green) at 4 dpi. Scale bar: 10 μm. **(F)** Quantification of the percentage of M1-macrophages at 4 dpi. Data of four replicates pooled. (Mean percentage/larva ± SEM, 4 dpi n_Sal_= 23 larvae, n_PBS_= 20, One sample Wilcoxon test, **** *P* < 0.0001).

To check macrophage polarization in persistently infected zebrafish, we analyzed infected Tg(*mfap4:mCherry-F*/*tnfa:GFP-F*) larvae at 4 dpi (**Fig. 5D**). Macrophage volume analysis revealed that few *tnfa*-positive macrophages (*mfap4^+^tnfa*^+^ cells) were still present at the infection site (less than 10%), but that the vast majority of accumulated macrophages were *tnfa*-negative (**Fig. 5D-E-F** and **Fig. S3B-C**). In agreement with the finding that the vast majority of accumulated macrophages at 4 dpi were not expressing *tnfa*, persistent *Salmonella* were found within *tnfa**-***negative macrophages at late stages of infection (**Fig. 5E**).

These results reveal that recruited macrophages switch their phenotype from M1-like states toward non-M1 states during the time course of *Salmonella* infection and that *Salmonella* survives in non-M1 macrophages at late stages of infection.

### Macrophages display dynamic transcriptional profiles upon *Salmonella* infection

To interrogate the molecular basis of macrophage activation during *Salmonella* infection, we compared the transcriptomic profiles of “activated” macrophages in infected host versus “inactivated” macrophages in non-infected condition at 4 hpi and 4 dpi. In our experimental setup, Tg(*mfap4:mCherry-F*/*tnfa:GFP-F*) larvae were either infected with *Salmonella* (*Sal*-INF) or not infected (Non-INF) (**Fig. 6A**). Fluorescence-Activated Cell Sorting (FACS) analysis on whole larvae in Non-INF condition revealed that a large majority of *mfap4*^+^ cells (mCherry^+^) were *tnfa*^-^ (GFP^-^) both at 4 hpi and 4 dpi (100% and 90.4% ± 4, respectively) (**Fig. S4A**). These expected populations were referred to as “inactivated” macrophages. FACS analysis in Sal-INF condition at 4 hpi revealed that the majority (92.5% ± 6) of *mfap4*^+^ cells (mCherry^+^) were *tnfa*^+^ (GFP^+^), while the majority (80.9% ± 4) were *tnfa*^-^ (GFP^-^) at 4 dpi (**Fig. S4A**). Therefore, *mfap4*^+^ *tnfa*^+^ cells at 4 hpi, which represent the main macrophage population of early infection phase, were referred to as “M1-activated”. In contrast the main macrophage population at 4 dpi consists of *mfap4*^+^ *tnfa*^-^ cells that were referred to as “non-M1-activated”.

**Fig. 6.**
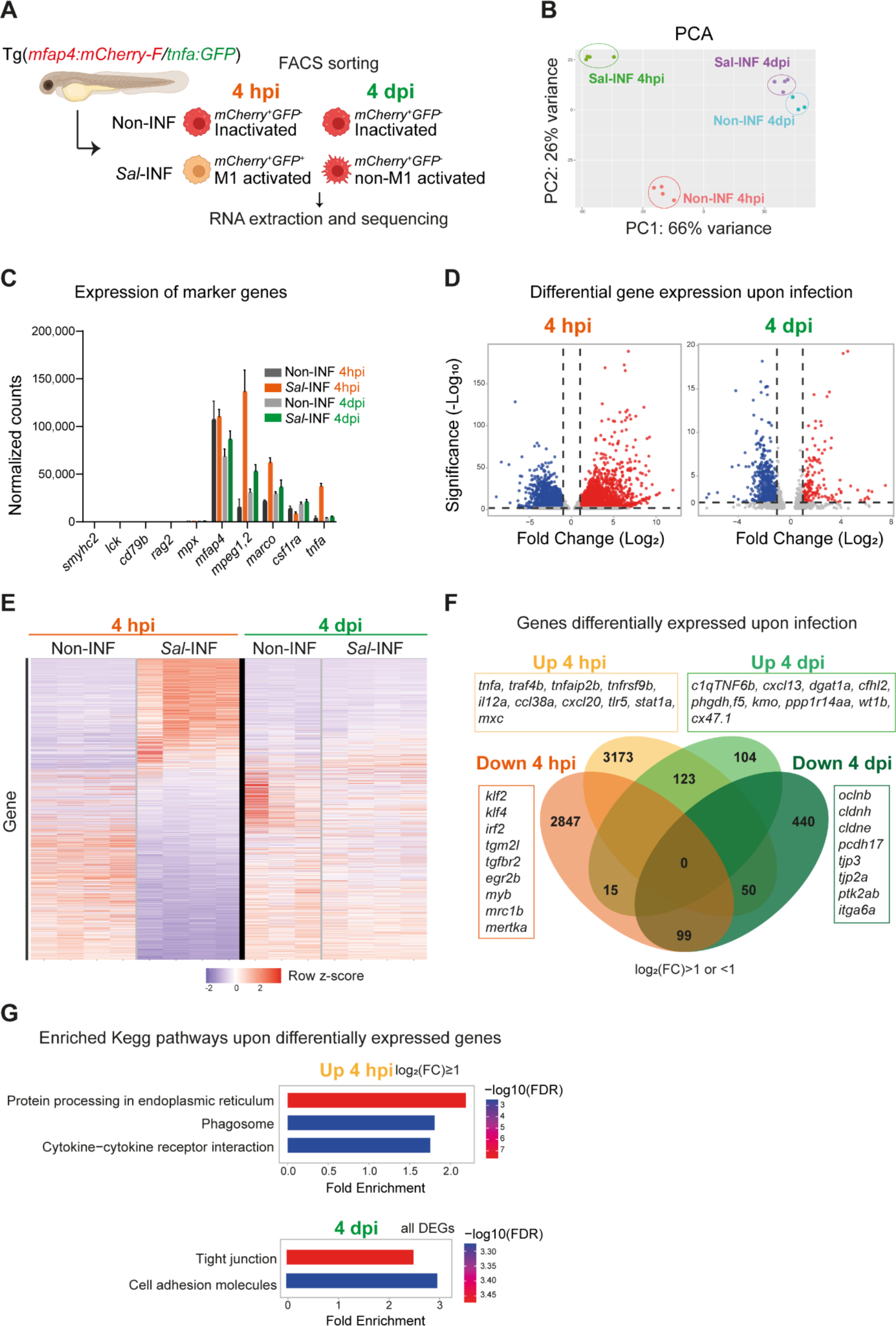
RNAseq analysis reveals macrophage transcriptome switch during *Salmonella* infection. **(A)** Schematic diagram of macrophage RNA-sequencing experimental design. Tg(*mfap4:mCherry-F*/*tnfa:GFP-F*) larvae were either infected with *Salmonella* (*Sal*-INF) or non-infected (Non-INF). FACS sorting was used to isolate *mfap4*^+^*-tnfa*^-^ cells (mCherry^+^ GFP^-^) and *mfap4*^+^*-tnfa*^+^ cells (mCherry^+^ GFP^+^) at 4 hpi and 4 dpi. **(B)** Principal Component Analysis (PCA) score plot of *mfap4*^+^*-tnfa*^-^ cells in Non-INF condition (n = 4) and *mfap4*^+^-*tnfa*^+^ cells in *Sal*-INF condition (n = 4) at 4 hpi and of *mfap4*^+^-*tnfa*^-^ cells in Non-INF condition (n = 3) and *mfap4*^+^-*tnfa*^-^ cells in *Sal*-INF condition (n = 4) at 4 dpi. **(C)** Normalized expression of several marker genes of muscle cells, lymphocytes, neutrophils and macrophages in the different sorted macrophage populations. **(D)** Volcano plot showing Differentially Expressed Genes (DEGs) between Non-INF and *Sal*-INF conditions at 4 hpi and 4 dpi. Adjusted p-value (*P*-adj) < 0.05 was used as the threshold to judge the significance of the difference in gene expression. Red plots: up-regulated genes; blue plots: down-regulated genes; grey plots: unchanged genes. **(E)** Heatmap of DEGs between macrophage populations across infection (*P*-adj < 0.05|Log_2_(FC)≥ 1). Selected top DEGs from each population are shown. Color-coding, decreased expression: blue, no expression: white, high expression: red. **(F)** Venn diagram showing unique and intersecting up-regulated or down-regulated genes (indicated as Up and Down, respectively) upon infection from macrophage transcriptome at 4 hpi and 4 dpi. The numbers of up-regulated and down-regulated gene are indicated in bold in each unique and overlapping sector of the Venn diagram. The most noteworthy genes of each unique sector of the Venn diagram are indicated (*P*-adj<0.05| (Log_2_ (FC)≥1 or ≤1)). **(G)** Chart representation of Kegg pathways enriched in up-regulated genes (*P*-adj< 0.05 | Log_2_ (FC)≥1) at 4 hpi (upper panel) and all DEGs at 4 dpi (lower panel) (*P*-adj < 0.05). Graph shows the fold enrichment, red color: lowest enrichment FDR and blue color: highest enrichment FDR.

Global gene expression analysis on these different populations was performed by RNA sequencing (**Fig. 6A**). The Principal Component Analysis (PCA) showed that each biological sample formed distinct clusters according to the 4 hpi and 4 dpi experimental conditions (**Fig. 6B**). As expected from the FACS procedure, all sorted populations expressed considerable levels of several key macrophage markers such as *mfap4*, *mpeg1.2*, *marco* and *csf1ra,* but not muscle marker (*smyhc2*), lymphocyte markers (*lck, rag2, cd79b*), neutrophil marker (*mpx*) neither other cell type markers (**Fig. 6C**). In addition, only the cells sorted as *mfap4*^+^ *tnfa*^+^ from the Sal-INF condition at 4 hpi expressed *tnfa* (**Fig. 6C**). To compare the functional differences between “inactivated” and “activated” macrophages, we performed a Differentially Expressed Gene (DEG) analysis between Non-INF and Sal-INF conditions at 4 hpi and 4 dpi (adjusted *p* value (*P*-adj) <0.05|log_2_(Fold Change (FC)) ≥ 1). *Salmonella* infection induced massive changes of gene expression in macrophages at 4 hpi (**Fig. 6D, Table S1**) and M1-activated macrophages harbored specific gene expression signature with 3,173 up-regulated and 2,847 down-regulated genes, whereas at 4 dpi, only 104 up-regulated and 440 down-regulated genes were specific to the non-M1 signature (**Fig. 6E-F, Fig. S4B** and **Table S1**). These results reveal that macrophages display a highly dynamic signature during the time course of infection.

At 4 hpi, M1-activated macrophages were characterized by a specific up-regulation of pro-inflammatory genes including those involved in the TNF pathway (*tnfa*, *traf4b*, *traf7*, *traf1b*, *tnfrsf1b, tnfrsf9b, tnfaip2* and *tnfaip3*), pro-inflammatory cytokines (*il12a*, *il12bb*), chemotaxis (*ccl38a.5, ccl38a.4, ccl19, cxcl18b, cxcl20*), complement (*c3a.1*, *c3a.3*, *c3a.4*, *c3a.6*), innate immune sensing of pathogens and antimicrobial responses (*tlr5*, *stat1a*, *stat2*, *mxc*, *irf9*, *crfb17*, *card11*, *card9, hif1ab*) and matrix metalloproteinases (*mmp9, mmp14a, mmp14b, mmp11b, mmp13b, mmp30, mmp23bb)* (**Fig. 6F**). Interestingly, *il10* and *il6*, that were described for their dual role during inflammation ^32, 33^, were also up-regulated, while known markers of M2 and anti-inflammation such as *mrc1b*, *tgm2l*, *tgfbr2, klf2, klf4, irf2, mertka* were down-regulated. Up-regulated genes at 4 hpi were classified according the gene ontology (GO) terms and the KEGG pathways (**Fig. S4C, Table S1**). Top ranking GO terms enriched in macrophages at 4 hpi were “inflammatory response”, “neutrophil migration”, “macrophage migration” and “defense response to gram negative bacterium”. Enriched KEGG pathways were “Protein processing in endoplasmic reticulum”, “phagosome” and “cytokine – cytokine receptor interactions” (**Fig. 6G, Table S1**). Together, these results confirm the pro-inflammatory signature of *tnfa*^+^ macrophages at 4 hpi.

By contrast non-M1-activated macrophages at 4 dpi specifically up-regulated genes involved in anti-inflammation and M2 markers in mammalian systems (*c1qtnf6b*, *cxcl13*, *dgat1a*, *cfhl2* (ortholog of *F13B* in human), *phgdh*, *f5, kmo* and *ppp1r14aa*), retinoic acid pathway (*rdh10a*, *rdh12*, *ugt1a*), bacterial defense *(npsn*, *c9*, *hamp and nos2b*), regeneration (*wt1b*), tissue protection and development (*cx47.1, lta, ecm1b*) and extracellular matrix remodeling (*mmp25b*) (**Fig. 6F, Table S1**). Interestingly, some pro-inflammatory genes, like *saa*, *cxcl11.1, s100b, ptgs2a, itgb3b, illr4* (*CLEC* orthologous) and *pppr2r2bb* were up-regulated at both times points but with a higher fold change at 4 hpi than 4 dpi, while others like *noxo1a*, *lsp1*, *plat*, *lpar1*, *selm*, *elf3* and *irf6* were up-regulated at 4 hpi and down-regulated at 4 dpi. By contrast, genes involved in anti-inflammatory response and tissue regeneration (*prg4a*, *cxcl19*, *cd59*, *clic2*) were down-regulated at 4 hpi and up-regulated at 4 dpi (**Fig. S4B**). Of note, some markers of microglia (*apoeb*, *ccl34b.1*, *csf1rb*) were down-regulated at both time points, suggesting a re-differentiation of this cell population upon HBV-infection. KEGG pathways analysis performed on DEGs at 4 dpi revealed the absence of a pro-inflammatory signature and that the most significantly enriched pathways were “tight junction” and “cell adhesion” (**Fig. 6G, Table S1**), suggesting profound changes in their polarization status and cell-adhesion program. Surprisingly, the most significantly enriched GO terms in up-regulated genes included “Astrocyte”, “Myelination”, “Oligodendrocyte” and “Myelin maintenance” (**Fig. S4D, Table S1**). Macrophages may express genes supporting astrocyte and oligodendrocyte function in the injured microenvironment of the infected HBV or these up-regulated genes may reflect the presence of internalized transcripts after efferocytosis in the infected area, as previously observed in other biological context ^34^. Altogether, these results reveal that macrophages skew their phenotype from acute to persistent infection, switching from a pro-inflammatory phenotype to an anti-inflammatory/pro-regenerative phenotype with a unique cell-adhesion signature.

### *Salmonella* persistence in the host is accompanied with macrophages harboring low motility

Our transcriptomic analysis data reveal that many genes related to cell adhesion are differentially regulated in *Salmonella* infection compared to control and that this cell adhesion signature is not the same during acute and persistent infections (**Fig. 7A**). These genes encompassed cell-cell adhesion molecules, likes cadherins (*cdh1*, *cdh2*, *cdh11, cdh17* and *cdh18*), occludins (*oclna*, *oclnb*, *cldnk*, *cln19*) and tight junction proteins (*tjp1a* (also known as zo-1), *tjp3*, *tjp2a*, *tjp2b*), extracellular matrix-adhesion proteins (*itgb1a*, *itgb1b*, *itgb4*, *itga6a*, *cd44a*), regulators of cell-adhesion (*pxna, ptk2ba, ptk2ab)* and regulators of cytoskeleton remodeling (*rac1a*, *rock2b, rhoua*) and most of them are down-regulated at 4 dpi (**Fig. 7A**). This shift in adhesion program of activated macrophages during the establishment of persistent infection suggests profound changes in their interaction with their environment and their migratory properties. Indeed, macrophage function relies on the establishment of contact with neighboring cells and stable attachments to their substrate, allowing transmigration through tissues, positioning and signaling. Cadherins were shown to regulate macrophage functions ^35, 36^ and integrins and small GTPases are also important for macrophage migration ^37, 38^. To test whether the changes in expression of cell-adhesion and cytoskeletal regulators were mirrored by changes in macrophage adhesion associated motility, Tg(*mfap4:mCherry-F*) larvae were infected with Sal-E2Crimson and imaged using confocal microscopy during 2 hours at 1 hpi and at 4 dpi. Analysis of macrophage trajectories at the injection site revealed that macrophage motility at 1 hpi was enhanced in infected larvae, with long-distance migration compared to PBS control (**Fig. 7B** and **Movies S8-9**). In contrast, macrophage motility at the injection site at 4 dpi in infected larvae was similar to that of PBS control, with very short trajectories. In addition, macrophage migration speed was significantly enhanced at 1 hpi in the infected larvae compared to the control condition, whereas at 4 dpi, this speed remained low and similar to that of the control condition (**Fig. 7C**). These data demonstrate that during acute infection, high motility macrophages are recruited to the infection site, while during persistent infection, clusters of motionless macrophages are formed while they provide a niche for *Salmonella* to survive.

**Fig. 7.**
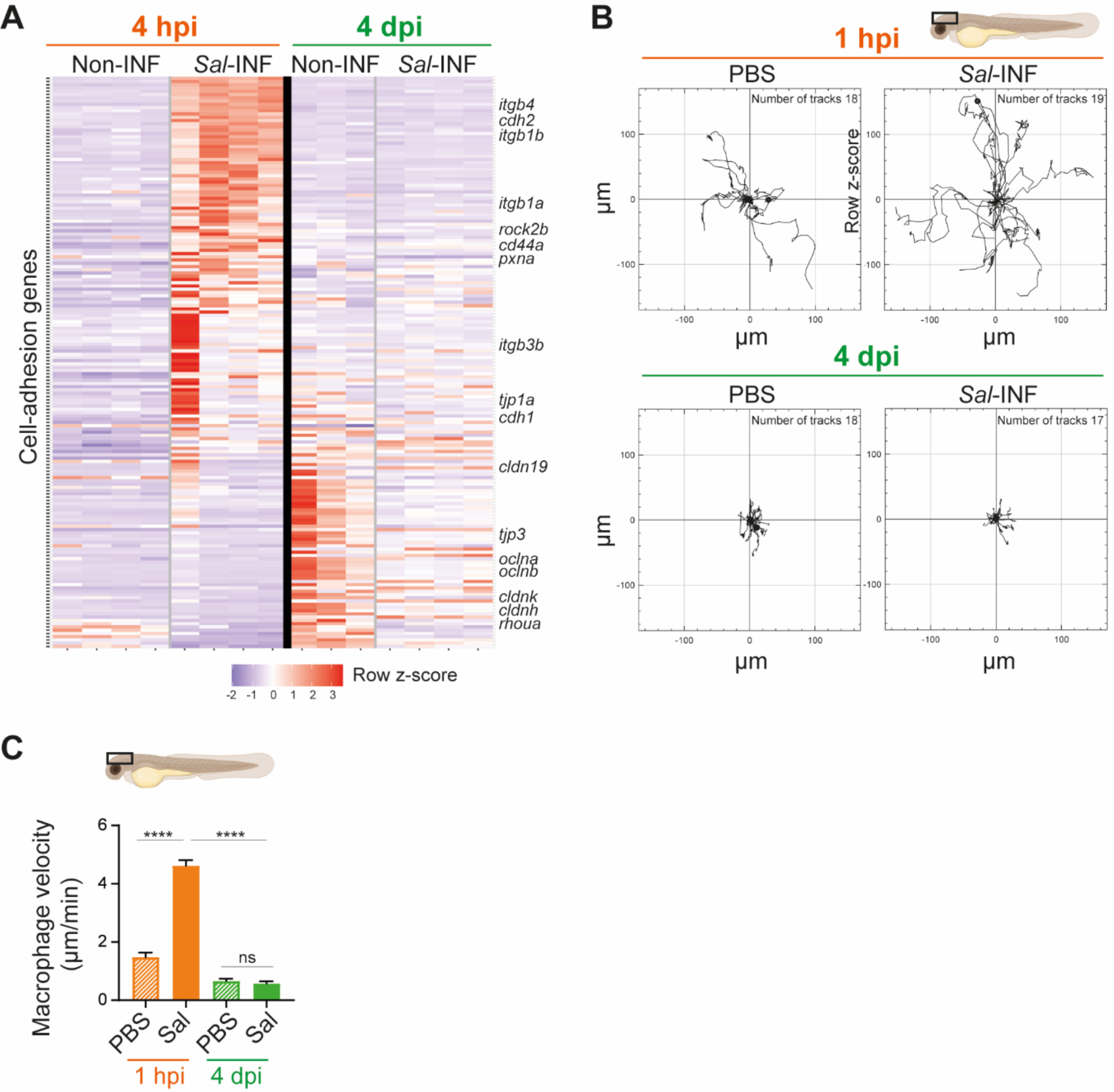
*Salmonella* persistence induces drastic changes in cell adhesion related-gene expression and motility in macrophages. **(A)** Heatmap of Differentially Expressed Genes (DEGs) involved in cell adhesion and tight junction, between macrophage populations across infection (*P*-adj < 0.05). Selected DEGs from each population are indicated. Color-coding, decreased expression: blue, no expression: white, high expression: red. **(B-C)** Tg(*mfap4:mCherry-F*) larvae were injected with PBS or Sal-E2Crimson in HBV and time-lapse videos of labelled macrophages were acquired during 2 hours at 1 hpi or 4 dpi. **(B)** Migration of macrophages in response to PBS or Sal-E2Crimson at 1 hpi or 4 dpi. Representative trajectory plots of individual macrophage movement tracks are shown, with the initial position in the center of the graph. Number of macrophage tracks are indicated. **(C)** Quantification of the individual macrophage velocity from PBS-injected or Sal-infected larvae at 1 hpi and 4 dpi. Data of two replicates per time point pooled. (Mean velocity/macrophage ± SEM, at 1 hpi: n_PBS_=26, n_Sal_=70; at 4dpi: n_PBS_= 52, n_Sal_=78; Mann-Whitney test, two-tailed, significance of Sal versus PBS conditions, **** *P* < 0.0001, ns: not significant; Anova test, significance of Sal-1hpi versus Sal-4dpi conditions).

## DISCUSSION

While some pathogens transiently infect an organism, others can survive inside the host for a long period of time or even for life. *Salmonella* can persist inside macrophages for long term ^3, 39^ and this persistence has been proposed to depend on macrophage polarization status ^5^. Here we describe the first model of *Salmonella* persistence in transparent zebrafish that allows the detailed analysis of the dynamics of interactions between *Salmonella* and polarized macrophages during early and late phases of infection in a whole organism.

In other zebrafish infection models with *Salmonella*, acute symptoms – hyper-proliferation of the bacteria, larval mortality – presumably prevent the establishment of a long-term infection ^18, 19^. Unlike these studies, we injected *Salmonella* in a closed compartment, the larval HBV. This mode of infection led to diverse outcomes, including acute infection, clearance and persistence. Indeed, in 47% of the larvae, *Salmonella* replicated intensively, leading to the rapid larval death. In surviving larvae, the bacteria were either cleared or survived inside the host, possibly up to 14 dpi. These diverse outcomes of infected zebrafish could be explained by multiple factors, including stochastic cell-to-cell differences in genetically identical bacteria or the complexity of immune cell population. Indeed, in *in vitro* systems, heterogeneous activity of the bacteria creates phenotypically diverse bacterial subpopulations which shape different cellular environments and potentiate adaptation to a new niche ^15, 40^.

We focused our *in vivo* study on embryos infected with persistent *Salmonella* up to 4 dpi. During the course of infection, the infected host displayed two main phases characterized by distinct inflammatory status: an early pro-inflammatory phase and a late phase characterized by both pro- and anti-inflammatory signals. We analyzed the interaction of *Salmonella* with innate immune cells during both phases. Similarly to previous work ^41^, during early infection, both neutrophils and macrophages engulfed the bacteria. Macrophages played a crucial role in bacterial clearance, as shown by the increased bacterial burden upon macrophage depletion. Exploiting the imaging possibilities of this system, we visualized macrophage activation thanks to *tnfa* expression in real time in response to *Salmonella* infection and showed that engulfing macrophages and bystanders polarized toward M1-like phenotypes. This pro-inflammatory response from the host was rapid and robust as more than 90% of the macrophages activated as M1-like within the first hours. Besides, we assessed the full transcriptome of zebrafish macrophages during early infection and we demonstrated that macrophages adopt a pro-inflammatory program characterized by the up-regulation of genes involved in inflammation, cytokine-cytokine receptor and phagosome pathways. These results are consistent with our initial analysis on whole larvae and are reminiscent to previous studies using whole *Salmonella*-infected larvae where genes encoding matrix metalloproteinases, pro-inflammatory cytokines and chemokine pathways are up-regulated ^18, 30^. There are also similarities with the activation program detected in human macrophages infected with different pathogens including *Salmonella* ^11^, which may represent an evolutionarily conserved program for host defense.

During the persistent phase, high resolution intra-vital imaging showed that *Salmonella* mainly localized inside macrophages that are organized in large clusters. By contrast, although some neutrophils are detected in the HBV, they are poorly mobilized, not clustered and do not harbor persistent bacteria. Importantly, most of the mobilized macrophages do not express *tnfa* in late stages of infection; we could not observe infected macrophage expressing *tnfa*. Transcriptomic profiling of these non-inflammatory macrophages during bacterial persistence revealed an anti-inflammatory and regenerative profile characterized 1/ by the attenuation of pro-inflammatory genes and 2/ by the up-regulation of genes involved in anti-inflammation, tissue maintenance, development and regeneration. Surprisingly, neuronal function-related genes, involved in myelination or expressed in astrocytes and oligodendrocytes were also up-regulated. This should not result from cell contamination given the macrophage-specificity of the RNAseq data sets, which did not recover other cell type markers. We therefore propose two possible scenarios. In the first scenario, the brain damages caused by the persistent infection may trigger a regenerative response involving pro-regenerative macrophages, which participate to the regeneration of axons and the restoration of their function by remyelination and maintenance of myelin homeostasis ^42–44^. Alternatively, during persistent infection, macrophages may perform efferocytosis of damaged or dying cells and internalize transcripts originating from them. The internalization of apoptotic RNAs by macrophages during efferocytosis has been demonstrated in the context of murine macrophages co-cultured with apoptotic human T cells ^34^. Both scenarios suggest that the macrophages interact tightly with their microenvironment during persistent infection.

Our data indicate that Non-M1 macrophages which have limited bactericidal activity, can be used by *Salmonella* as a niche to sustain persistent infections in their hosts. *Salmonella* persistence was previously associated with M2 macrophages in a mouse model of long-term infection ^5, 14^. Molecularly, M2-permissive macrophage polarization in granulomas is partially dependent on the bacterial factor SteE, while the host cytokine TNF limits M2 polarization ^13^. In infected cultured-macrophages, SteE regulates STAT3, reorienting macrophage polarization towards M2 phenotypes ^45, 46^. Another study revealed that macrophages harboring *Salmonella* Typhimurium during persistent stages express high levels of the inducible nitric oxide synthase (known as iNOS or NOS2) ^14^. Interestingly, we also observed an up-regulation of *nos2b* in zebrafish macrophages four days after *Salmonella* infection. Furthermore, non-growing, antibiotic-tolerant bacteria, also called persisters, were shown to translocate *Salmonella* pathogenicity island 2 type III secretion system effectors into the macrophage to reprogram it into a non-inflammatory macrophage ^47^. Our data are in line with these studies, emphasizing the potential of the zebrafish model for the study of persistent infections. However, further investigations are still needed to identify the bacterial factors involved in macrophage reprogramming in our system.

An efficient immune response requires the interplay of a cocktail of cell-adhesion molecules and immune cells ^48, 49^. Macrophages express various cell-adhesion proteins including *integrin b1* (*itgb1*) that is crucial for cell movements and protrusiveness during surveillance ^38^. Tight junctions are important for particle uptake and exchange ^50^ and RAC proteins were shown to be necessary for macrophage basic motility and migration ^51, 52^. We demonstrated a shift in macrophage adhesion program during persistent stages with several cell adhesion-related genes that were down-regulated, including *itgb1*, *rac2* and tight junction-related genes. This shift in adhesion program was accompanied with a decrease in macrophage motility. As these motionless macrophages appear non-inflammatory and contain persistent bacteria, they may constitute a permissive environment for bacterial survival. Subversion of macrophage motility has been previously observed with another pathogen, *Mycobacterium marinum*. In this case, mycobacteria use the ESX-1/RD1 secretion system to enhance macrophage recruitment to nascent granulomas and favor granulomas formation ^53^.

Granulomas are a key pathological feature of some intracellular bacterial infections; they do contain a high proportion of macrophages. *Salmonella enterica* can cause granuloma formation in different animal species ^54^, including mice ^14^ and humans ^55–57^. The complex cellular structure of granuloma is thought to be important for bacterial persistence. Recently, using single cell transcriptomics, Pham and collaborators identified divers macrophage populations within *Salmonella* Typhimurium induced granulomas ^58^. In this study, *Salmonella* persisted within aggregates of *tnfa*-negative macrophages that remind early granulomas. Mycobacterial granulomas are characterized by epithelioid macrophages expressing tight junction proteins, E-cadherin (CDH1) and ZO-1 (*tjp1)* ^59, 60^. Interestingly, in the context of persistent *Salmonella* infection, we showed that *tnfa*-negative macrophages down-regulated the expression of several tight-junction related genes during persistence, such as *cdh1* and *tjp1a*. This is similar to what has been observed in *Salmonella*-induced granulomas in mice where macrophages weakly express E-cadherin or ZO-1 ^14^ and suggests that these structures are non-epithelioid granulomas initiated by an innate immune mechanism.

In conclusion, the zebrafish larva proves to be an extraordinary platform for the analysis of *Salmonella* persistent infection and the understanding of long-term interactions with host cells inside a whole living animal. The highly dynamic changes of macrophages gene expression between early infection and persistent phases confirm the highly versatile nature of macrophage-*Salmonella* interactions and suggest that macrophage polarization and motility switch play an important role in the establishment of a secure niche for *Salmonella* (**Fig. 8**).

**Fig. 8.**
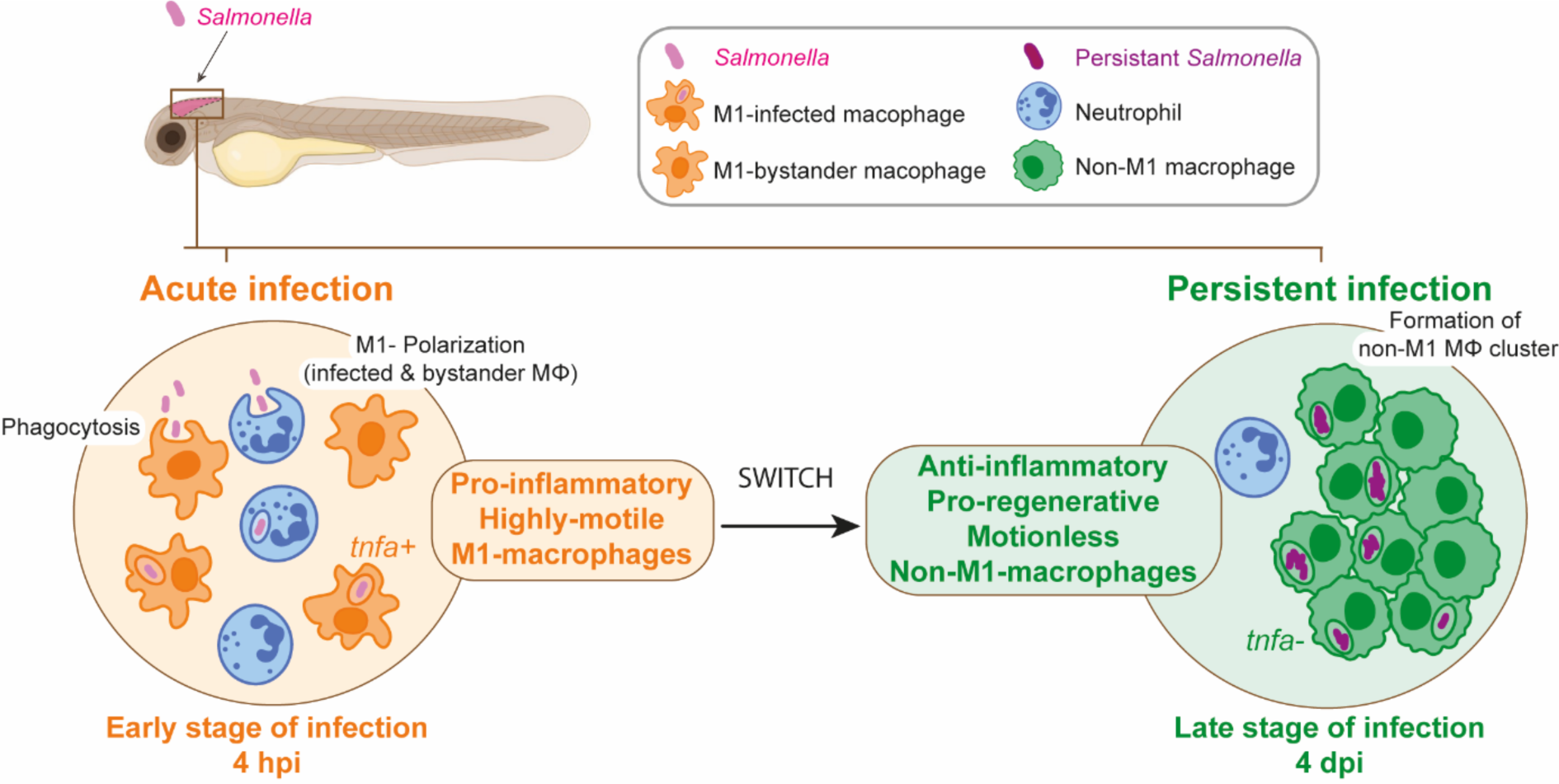
From acute to persistent *Salmonella* infection, macrophages switch their polarization states and motility. Schematic representation of the two main phases of *Salmonella* HBV-infection in zebrafish. The early phase corresponds to an acute infection characterized by the recruitment of leukocyte populations, phagocytosis of *Salmonella* and M1-polarization of highly motile macrophages, while during the late phase *Salmonella* persists inside motionless macrophages that display an anti-inflammatory and pro-regenerative status and form clusters.

## MATERIALS AND METHODS

### Ethics statement

Animal experimentation procedures were carried out according to the European Union guidelines for handling of laboratory animals (https://ec.europa.eu/environment/chemicals/lab_animals/index_en.htm) and were approved by the Comité d’Ethique pour l’Expérimentation Animale under reference CEEA-LR-B4-172-37 and APAFIS #36309-2022040114222432 V2. Fish husbandry and embryo collection and experiments were performed at the University of Montpellier, LPHI/CNRS UMR5235.

### Fish husbandry

Fish maintenance, staging and husbandry were performed as described ^61^ with golden, AB strains and transgenic lines. *Tg(mfap4:mCherry-F)ump6tg* ^62^ is here referred to as *Tg(mfap4:mCherry-F). Tg(mpx:eGFP)ill4* referred to as *Tg(mpx:eGFP)* ^63^ was used to visualize neutrophils. *Tg(tnfa:GFP-F)ump5Tg* referred to as *Tg(tnfa:GFP-F)* was used to visualize cell expressing *tnfa* ^17^. *Tg(mfap4:mCherry-F)* crossed with *tg(tnfa:GFP-F)* were used to visualize activated M1 macrophages. *Tg(mpeg1:Gal4FF)gl25* ^64^ and *Tg(UAS-E1b:nfsB-mCherry)i149* ^65^ were used to ablate macrophages. Embryos were obtained from pairs of adult fishes by natural spawning and raised at 28°C in tank water. Embryos and larvae were staged according to ^66^ and used for experiments from 0 hpi to 17 dpi. Larvae were anaesthetized in zebrafish water supplemented with 200 μg/ml tricaine (Ethyl 3-aminobenzoate methanesulfonate, MS-222 Sigma #A5040) before any manipulation (infection or imaging) and if necessary were replaced in their medium at 28°C.

### Salmonella strains

*Salmonella* strains were grown overnight in Luria-Bertani (LB) medium at 37°C with 100 µg/ml ampicillin, 10 µg/ml tetracycline or 25 µg/ml kanamycin when required. *Salmonella enterica* serovar Typhimurium ATCC14028s (here called *Salmonella*) was used as the original parental *Salmonella* strain. *Salmonell*a carrying plasmid pRZT3::dsRED ^19^, pE2-Crimson (Clontech) and pFPV25.1 ^67^, that express red fluorescent protein (dsRED), far-red fluorescent protein (E2Crimson) and green fluorescent protein (GFP) respectively, were used for micro-injection in zebrafish embryos (see below). To create a *Salmonella* strain expressing chromosomal copies of GFP, the *rpsM::gfp* fusion from strain SM022 ^68^ was transferred by P22 transduction of the *rpsM::gfp* fusion, linked to kanamycin resistance gene into the original parental *Salmonella* strain ATCC14028s through selection for kanamycin resistance.

### Salmonella injections

*Salmonella* strains were grown to exponential phase and recovered by centrifugation, washed twice and resuspended in PBS at an OD_600_ of 5 or 2.5 (depending on the required dose) with phenol red. Infection was carried out by microinjection of 1.5 nL of bacterial suspensions in the Hindbrain compartment of dechorionated and anesthetized 2 dpf embryos. Two different doses of *Salmonella* were used for microinjection in zebrafish embryos: low (<500 CFU) and high (1000-2000 CFU). The inoculum dose was checked by counting the CFU containing in 1.5 nL of the bacterial suspension. Per replicate a minimum of 30 larvae were infected and three replicates were done for each experiment.

### Quantification of bacterial load by CFU counts

A minimum of five larvae per time points were anaesthetized in zebrafish water supplemented with 200 μg/ml tricaine and then each embryo was crushed in 1% Triton-X100-PBS in an Eppendorf tube using a pestle (MG Scientific #T409-12). After 10 min incubation at room temperature, dilutions of total lysates were plated on LB agar plates containing appropriate antibiotics. CFU were counted after an overnight incubation of the plates at 37°C. Larvae used for CFU counts were randomly chosen among surviving larvae.

### Macrophage ablation

For macrophage depletion, we used Tg(*mpeg1:Gal4/UAS:nfsB-mCherry*) embryos expressing *nfsB-mCherr*y under the indirect control of *mpeg1* promoter. *nfsB-mCherry* encodes an *Escherichia coli* nitroreductase (NTR) fusionned to mCherry protein that converts metronidazole (MTZ) into a toxic agent that kills cells. Tg(*mpeg1:Gal4/UAS:nfsB-mCherry*) embryos were incubated in zebrafish water containing 10 mM Metronidazole (MTZ, Sigma-Aldrich) and 0.1% DMSO at 48 hpf and 24 h before injection with *Salmonella* or PBS. Treatment with 0.1% DMSO was used as a control. Depletion efficiently was assessed by imaging using the MVX10 Olympus microscope just before HBV-injection. Effects of macrophage depletion on embryo survival and bacterial load during infection were analyzed at 1, 2, 3 and 4 dpi.

### Imaging of live zebrafish larvae

Larvae were anesthetized in 200 μg/ml tricaine, positioned on 35-mm glass-bottomed dishes (WillCo-dish®), immobilized in 1% low-melting-point agarose and covered with 2 ml of embryo water supplemented with 160 μg/ml tricaine. Epi-fluorescence microscopy was performed by using an MVX10 Olympus MacroView microscope that was equipped with MVPLAPO 1× objective and XC50 camera. Confocal microscopy was performed using an ANDOR CSU-W1 confocal spinning disk on an inverted NIKON microscope (Ti Eclipse) with ANDOR Neo sCMOS camera (20x air/NA 0.75 objective). Image stacks for time-lapse acquisitions were performed at 28°C. The 4D files generated by the time-lapse acquisitions were processed using ImageJ as described below. Three-dimensional reconstructions were performed on the four-dimension files for time-lapse acquisitions or on three-dimension files using Imaris (Bitplan AG, Zurich, Switzerland). In Fig. 3C and in Fig. S3A, a custom-made LSFM developed at ICFO was used ^69^. For the described LSFM experiments we made use of two illumination air objectives (Nikon 4x/NA 0.13), one water-immersion detection objective (Olympus 20x/NA 0.5), a sCMOS camera (Hamamatsu Orca Flash4.v2), and a 200mm tube lens (Thorlabs), obtaining an overall magnification of 22.2x. For LSFM imaging, zebrafish embryos were embedded within a Fluorinated Ethylene Propylene (FEP) tube (ID 2 mm, OD 3 mm) containing 0.2% LMP agarose with the addition of 160 μg/ml tricaine. The inner walls of the FEP tube would have been previously coated with a 3% methyl cellulose layer to avoid tail adhesion. After plugging with 1.5% LMP agarose the bottom end of the FEP tube, it was inserted into the LSFM imaging chamber and mounted vertically and upside-down. The temperature-controlled chamber was then filled with 15 mL of embryo water supplemented with 160 μg /ml tricaine.

### Visualization of interaction between *Salmonella*, macrophages and neutrophils

The 3D and 4D files generated by confocal microscopy were processed using ImageJ. First, stack of images from multiple time points were concatenated and then brightness and contrast was adjusted for better visualization with the same brightness and contrast per channel for every infected and PBS-control larva in each experiment. To generate the figure panels, stacks of images were compressed into maximum intensity projections and cropped. For visualization of bacteria localization related to macrophages and neutrophils, surfaces tool of Imaris software was used to reconstruct the 3D surfaces of leukocytes and bacteria.

### Quantification of total leukocyte population, quantification of recruited neutrophils and macrophages

To quantify total leukocyte populations, transgenic reporters were tricaine-anesthetized and whole larvae were imaged using MVX10 Olympus microscope. Total numbers of fluorescent leukocytes were counted by computation using Fiji (ImageJ software) as following: 1/ leukocytes (Leukocyte Units, LU) were detected using “Find Maxima” function, 2/ Maxima were automatically counted using run (“ROI Manager . . .”), roi-Manager(“Add”) and 3/ roiManager(“Measure”) functions. To quantify recruited leukocyte populations in the HBV, only the infection sites of reporter larvae were imaged using Spinning disc Nikon Ti Andor CSU-W1 microscope. ImageJ was used to concatenate stacks of images containing multiple time points and to adjust and set the same brightness and contrast per channel for every infected and PBS-control larva in each experiment. “Surfaces” tool of Imaris was used to reconstruct in 3D cell surfaces. Total volumes of immune cells were then extracted for relative quantification and expressed as mean volume (um^3^) with standard error of the mean. For the quantification of the percentage of M1-macrophages, the ratio of the total volume of *mfap4*^+^-*tnfa*^+^ cells among *mfap4*^+^ cells was calculated.

### RNA preparation on whole larva and quantitative RT-PCR analysis

For quantitative RT-PCR analysis, 2 dpf larvae were either injected in the HBV with Sal-GFP or with PBS as described above. To determine the relative expression of *il1b*, *tnfb*, *tnfa*, *il8*, *ccr2*, *ccl38a.4*, *mmp9*, *mrc1b*, *nrros*, *mfap4* and *mpeg1*, total RNA from infected larvae or controls (pools of 8 larvae each) was prepared at 3 hpi, 1 dpi, 2 dpi, 3 dpi and 4 dpi using Macherey-Nagel Nucleospin RNA Kit (# 740955.250). RNAs were reverse transcribed using OligodT and M-MLV reverse transcriptase (# 28025-013) according to manufacturer recommendations. Quantitative-RT-PCR were performed using LC480 and SYBER Green (Meridian BIOSCIENCE®, SensiFAST^TM^ SYBR # BIO-98050) according to manufacturer recommendations and analyzed using LC480 software. The final results are displayed as the fold change of target gene expression in infected condition relative to PBS control condition, normalized to *ef1a* as reference gene (mean values from 8 independent experiments with standard error of mean). The primers used are listed below.

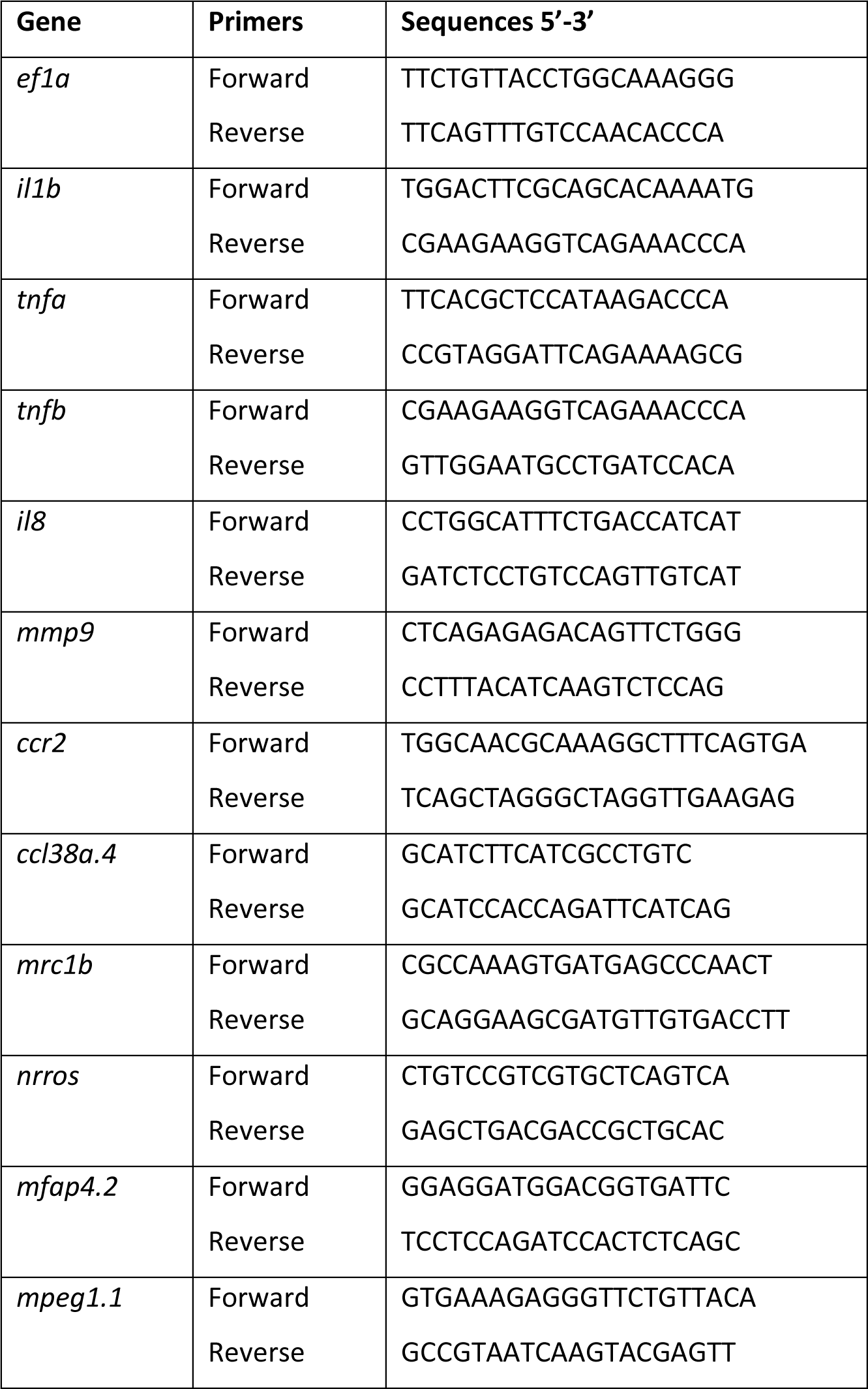

### Transcriptomic analysis on FACS-sorted macrophages

Double transgenic larvae, *Tg(mfap4:mCherry-F; tnfa:GFP-F)*, were either non infected (uninfected groups) or infected with non-labelled *Salmonella* at 48 hpf (infected groups), as described above. Cell dissociation from pools of 300 larvae were performed at 4 hpi and 4 dpi. At 4 dpi, only larvae harboring macrophage clusters in the brain were kept. Cell dissociation, FACS sorting and RNA preparation were performed as described in ^70^. A total of 15 samples were processed for transcriptome analysis using cDNA sequencing. The four experimental groups are: “uninfected/4 hpi”, “*Salmonella* infected/4 hpi”, “uninfected/4 dpi”, “*Salmonella* infected/4 dpi”. Experimental groups were obtained from four replicates, except for the condition “uninfected/4 dpi” which was obtained from three replicates. The 15 samples were sent to Montpellier GenomiX plateform (MGX, Institut de Genomique Fonctionnelle, Montpellier, France) for library preparation and sequencing. RNA-seq libraries were generated from 5 ng of RNA with the SMART-Seq v4 Ultra Low Input RNA Kit from Takara Bio (#634889) and the DNA Prep Kit from Illumina (#20060060) and were clustered and sequenced using an Illumina NovaSeq 6000 instrument, a flow cell SP and NovaSeq Xp Workflow according to the manufacturer’s instructions, with a read length of 100 nucleotides. Image analysis and base calling were done using the Illumina NovaSeq Control Software and Illumina RTA software. Demultiplexing and trimming were performed using Illumina’s conversion software (bcl2fastq 2.20). The quality of the raw data was assessed using FastQC (v11.9) from the Babraham Institute and the Illumina software SAV (Sequencing Analysis Viewer). FastqScreen (v0.15) was used to identify potential contamination. The RNAseq data were mapped on the zebrafish genome (version GRCz11) and gene counting was performed with Featurecounts v2.0.3. Sequencing depth of all sample was between 53 and 87 million reads. All reads were aligned to the GRCz11 version of the zebrafish genome using TopHat2 v2.1.1 (using Bowtie v2.3.5.1), software and Samtools (v1.9) was used to sort and index the alignment files. Subsequently, normalization and differential gene expression analysis was performed using DESeq2 and edgeR methods. After comparison of the two methods, DESeq2 method ^71^ was kept. Within DESeq2 p-values were adjusted using Benjamini & Hochberg (1995) corrections for controlling false positive rate (False Discovery Rate_FDR, also called p-adjusted (*P*-adj)) and results were considered statistically significant when *P*-adj< 0.05. Analysis was performed based on the log_2_ fold change (FC). Gene Ontology analysis (GO) of differentially expressed genes and KEGG pathways enrichment analysis were performed with ShinyGO 0.76 ^72^ with criteria of *P*.adj < 0.05 up-regulated ǀ log_2_(FC) > 0 or ≥ 1 and downregulated ǀ log_2_(FC) < 0 or ≤ 1.

The raw data is available in the NCBI GEO database under accession number: GSE224985, https://www.ncbi.nlm.nih.gov/geo/query/acc.cgi?acc=GSE224985

### Statistical analysis

Studies were designed to generate experimental groups of approximatively equal size, using randomization and blinded analysis. The sample size estimation and the power of the statistical test were computed using GPower. A preliminary analysis was used to determine the necessary sample size N of a test given α<0.05, power =1-β> 0.80 (where α is the probability of incorrectly rejecting H0 when is in fact true and β is the probability of incorrectly retaining H0 when it is in fact false). Then the effect size was determined. Groups include the number of independent values, and the statistical analysis was done using these independent values. The number of independent experiments (biological replicates) is indicated in the figure legends when applicable. The level of probability P < 0.05 constitute the threshold for statistical significance for determining whether groups differ and this p value is not varied throughout the Results. Graph Pad Prism 7 Software (San Diego, CA, USA) was used to construct graphs and analyze data in all figures. Specific statistical tests were used to evaluate the significance of differences between groups (the test and p value is indicated in the figure legend).

## ACKNOWLEDGEMENTS

This work was undertaken with support from the fish facility of Zefix-LPHI (University of Montpellier), Catherine Gonzalez (LPHI, CNRS) and the qPCR Haut Debit platform of the University of Montpellier. We acknowledge the imaging facility BioCampus Montpellier Ressources Imagerie (MRI), member of the national infrastructure France-BioImaging supported by the French National Research Agency (ANR-10-INSB-04, “Investments for the future”). We thank Dr. Ferric C. Fang (University of Washington School of Medicine) for *Salmonella* strain SM022. Dr Laurent Manchon for his advices for the RNAseq analysis. ICFO authors also acknowledge financial support from the Spanish Ministry of Economy and Competitiveness through the “Severo Ochoa” program for Centres of Excellence in R&D (CEX2019-000910-S), from Fundacio Privada Cellex, Fundacio Mir-Puig, Generalitat de Catalunya through the CERCA program, and Laserlab-Europe EU-H2020 (871124). MGX acknowledges financial support from France Génomique National infrastructure, funded as part of “Investissement d’Avenir” program managed by Agence Nationale pour la Recherche (contract ANR-10-INBS-09).

## AUTHOR CONTRIBUTIONS

J.L. & M.N-C. conceived experiments with input from A.B-P and G.L.. J.L., C.B-P., T.S., S.T., M.B., E.G., X.M., A.A.G. and L. B. performed experiments. J.L. & M.N-C. wrote the manuscript with input from A.B-P, G.L., T.S., M.B., E.G., P.L-A. and L. B.. M.N-C., G.L. and P.L-A. secured funding. All authors approved the manuscript.

